# Connectivity of the neuronal network for contextual fear memory is disrupted in a mouse model of third-trimester binge-like ethanol exposure

**DOI:** 10.1101/2024.08.16.608336

**Authors:** Mitchell D. Morningstar, Katalina M. Lopez, Stefanie S. Mayfield, Roberto N. Almeida-Mancero, Joshua Marquez, Andres M. Flores, Brooke R. Hafer, Edilberto Estrada, Gwen A. Holtzman, Emerald V. Goranson, Natalie M. Reid, Abigale R. Aldrich, Desna V. Ghatalia, Juhee R. Patel, Christopher M. Padilla, Glenna J. Chavez, Javier Kelly-Roman, Pooja A. Bhakta, C. Fernando Valenzuela, David N. Linsenbardt

**Author notes:** co-equal first authors (Mitchell D. Morningstar, Katalina M. Lopez). <u>Corresponding Authors:</u> David Linsenbardt, Ph.D. & C. Fernando Valenzuela, MD, PhD, Department of Neurosciences, MSC08 4740, 1 University of New Mexico, Albuquerque, NM 87131-0001 U.S.A.

## Abstract

**Background:** In rodents, third-trimester equivalent alcohol exposure (TTAE) produces significant deficits in hippocampal-dependent memory processes such as contextual fear conditioning (CFC). The present study sought to characterize changes in both behavior and Fos+ neurons following CFC in ethanol (EtOH)-treated versus saline-treated mice using TRAP2:Ai14 mice that permanently label Fos+ neurons following a tamoxifen injection. We hypothesized that TTAE would produce long-lasting disruptions to the networks engaged following CFC with a particular emphasis on the limbic memory system.

**Methods:** On postnatal day 7, mice received either two injections of saline or 2.5 g/kg EtOH spaced 2 hours apart. The mice were left undisturbed until they reached adulthood, at which point they underwent CFC. After context exposure on day 2, mice received a tamoxifen injection. Brain tissue was harvested. Slides were automatically imaged using a Zeiss AxioScanner. Manual counts on *a priori* regions of interest were conducted. Automated counts were performed on the whole brain using the QUINT 2D stitching pipeline. Last, novel network analyses were applied to identify future regions of interest.

**Results:** TTAE reduced context recall on day 2 of CFC. Fos+ neural density increased in the CA1 and CA3. Fos+ counts were reduced in the anteroventral (AV) and anterodorsal thalamus. The limbic memory system showed significant hyperconnectivity in male TTAE mice and the AV shifted affinity towards hippocampal subregions. Last, novel regions such as a subparafascicular area and basomedial amygdalar nucleus were implicated as important mediators.

**Discussion:** These results suggest that CFC is mediated by the limbic memory system and is disrupted following TTAE. Given the increase in CA1 and CA3 activity, a potential hypothesis is that TTAE causes disruptions to memory encoding following day 1 conditioning. Future studies will aim to determine whether this disruption specifically affects the encoding or retrieval of fear memories.

## Introduction

Fetal exposure to alcohol remains a significant health concern. Fetal Alcohol Spectrum Disorders (FASDs) is an umbrella term used to describe the consequences of this exposure. FASDs are the primary teratogen-induced cause of intellectual disabilities, leading to impairments in cognitive abilities that last until adulthood (Popova et al., 2023). First and second-trimester drinking has been of interest in most FASD studies. However, third-trimester drinking is still common in some regions of the United States where a prevalence of 8% for late pregnancy ethanol exposure has been reported (Umer et al., 2020). Similarly, a study conducted from 1997-2002 reported that 7.9% of pregnant women drink during this point in gestation (Ethen et al., 2009). Although many key developmental stages are nearing completion late in pregnancy, third-trimester-equivalent alcohol exposure (TTAE) can still have serious effects on fetal health. This is in part because the third trimester encompasses a key brain growth spurt that involves myelination, early synaptic development, and growth, which are particularly sensitive to ethanol exposure. While this growth spurt takes place during the third trimester in humans, it happens during the first two weeks of life in rodents (Dobbing & Sands, 1979).

Various paradigms have been used to study the effect of ethanol exposure during the brain growth spurt, including vapor chamber exposure, subcutaneous injections, and intragastric intubation. Subcutaneous ethanol injection during the brain growth spurt leads to significant apoptosis in rodents and rhesus macaques (Farber et al., 2010; Livy et al., 2003; Olney et al., 2002; Tenkova et al., 2003; Young & Olney, 2006). A widely used “double-hit” model consisting of two, 2.5 g/kg injections at postnatal day 7 has been shown to produce apoptotic neurodegeneration in the rodent hippocampus, striatum, dorsal thalamus, and cerebral cortex, all of which are a part of the limbic memory system (Ikonomidou et al., 2000; Saito et al., 2007). Additionally, the model produces long-lasting effects, including significant neuronal loss at post-natal day 70 (P70) in these brain regions (Smiley et al., 2023). However, the consequences of this insult on learning and memory remain relatively unknown. A study conducted two decades ago reported notable damage in spatial memory at one month of age in mice after confirming apoptosis and long-term neuronal loss in diencephalic structures (D. Wozniak et al., 2004).

Contextual fear conditioning (CFC) refers to the learning processes underlying the paired acquisition of a noxious stimulus, such as a foot shock, with a specific environment. The environment subsequently produces an unconditioned response (i.e. freezing) in the absence of the paired noxious stimulus. This allows for rapid and easily assessable fear-learning responses with minimal training necessary. Critically, TTAE produces deficits in CFC conditioning in rodents across multiple exposure and conditioning paradigms (Goodfellow & Lindquist, 2014; Hamilton et al., 2011; Heroux et al., 2019; Hunt et al., 2009; Wilson et al., 2016). However, the mechanisms underlying these TTAE-induced contextual learning deficits are not fully understood. This is partly because the contextual memory associated with the harmful event encompasses spatial and non-spatial attributes such as sounds, smells, lighting, and internal states (Marks et al., 2022; Vasudevan et al., 2024). Thus, contextual memories are necessarily encoded by neuronal networks located in multiple brain regions, including the limbic memory system. We, therefore, hypothesized that TTAE induces contextual memory deficits by disrupting the *connectivity* of these disparate brain regions, which we tested using recent advancements in neuronal labeling methods.

Tamoxifen-inducible Targeted Recombination in Active Populations, version 2 (TRAP2) mice were utilized to capture Fos activity during memory retrieval (DeNardo et al., 2019). Fos is a marker of neural activity encoded by the immediate early gene *c-fos* after intracellular calcium concentrations significantly exceed basal levels. Consequently, it is interpreted as a signal for heightened neural activation (Chung, 2015). Utilizing TRAP2 mice, we sought to determine if brain regions critical for the expression of CFC were altered following TTAE. These include areas such as the retrosplenial cortex (RSP) (Todd et al., 2019), hippocampal formation (McNish et al., 2000), and anterior thalamic nuclei (ATN) (Marchand et al., 2013). Next, utilizing a recently developed novel pipeline for histological processing named QUINT (Yates et al., 2019), we determined the effect of TTAE on functional Fos-labeled networks across the brain.

Functional Fos network analyses have recently gained traction in alcohol research (Ardinger et al., 2024; Kimbrough et al., 2020; Roland et al., 2023) as they can allow for unbiased discovery of brain regions critically involved in a behavior or affected by a treatment. Towards this, we sought to determine the brain regions affected by TTAE during contextual fear learning and additionally how their functional connectivity may be altered to other brain regions.

## Methods

### Ethanol exposure paradigm

All animal procedures were approved by UNM-HSC Animal Care and Use Committee and adhered to NIH guidelines. STOCK Fos^tm2.1(icre/ERT2)Luo^/J (*TRAP2*; stock #030323) and *B6.Cg-Gt(ROSA)26Sor^tm14(CAG-tdTomato)Hze^/J* (*Ai14;* stock #007914) mice were purchased from Jackson Labs. Female *TRAP2* mice were crossed to male *Ai14* mice to obtain the double heterozygous (*TRAP2;Ai14*) mice. Genotyping was performed by Transnetyx (Memphis, TN). Mice were maintained on a reverse light cycle (lights on at 8 pm) and had access to food (Teklad 2920x diet, Envigo, Indianapolis, IN) and water *ad libitum*. Mice were housed in a micro-isolated system with ventilated racks and cages (Lab products, Seaford, DE). On postnatal day 7, mice were injected with saline or ethanol (20% v/v in normal saline) subcutaneously (2 x 2.5 g/kg doses at a 2 h interval between doses). This paradigm produces peak blood ethanol concentrations between 400-500 mg/dl (D. Wozniak et al., 2004). A total of 49 litters were generated for the present set of experiments. After weaning, mice were housed in same-sex groups and allowed to mature until adulthood (3-4 months old) prior to behavioral studies. In the behavioral analyses and manual FOS count analyses, data from 37 Saline Females, 44 Saline Males, 40 TTAE Females, and 38 TTAE Males were evaluated. QUINT counts and functional connectivity analyses utilized a subset of data: 26 Saline Females, 18 Saline Males, 27 TTAE Females, and 25 TTAE Males.

### Contextual Fear Conditioning

We used the TRAP technique to fluorescently tag active neurons during contextual fear memory retrieval (DeNardo et al., 2019). This approach utilizes an immediate early gene promoter (c-Fos) to control the expression of a tamoxifen-inducible Cre-estrogen receptor (Cre-ER) construct that induces expression of a Cre-dependent effector (expression of the td-Tomato protein in our case). Activation of neurons induces expression of Cre-ER in the cytoplasm and administration of tamoxifen enables it to translocate into the nucleus where it causes recombination and permanent expression of td-Tomato in activated neurons. TRAP2;Ai14 mice were allowed to acclimatize to the experimental room for at least 30 min and were handled for 2 min/day for 5 days before conditioning. Mice were transported to and from the experimental room in their home cages using a wheeled cart. The mice were kept in an adjacent area prior to contextual fear experiments. The fear-conditioning chamber consisted of a square cage (20cm x 16cm x 25.5cm) with a grid floor wired to a shock generator (MedAssociates, #ENV-4145), surrounded by a sound insulation chamber (MedAssociates, #VFC-022MD). The chamber was illuminated with dim white and near-infrared lighting (MedAssociates, #NIR100) and was scented with 0.01% acetic acid. Day 1 of the fear conditioning protocol consisted of 4 shocks (2 s; 0.7mA) with an inter-shock interval of 2 minutes (total paradigm duration = 540 s; shocks delivered at 148, 268, 388, and 508 s). Twenty-four hours later (Day 2), the animals were placed in the chambers under the same conditions except that shocks were not delivered (total paradigm duration = 720 s). Freezing time during day 2 was determined using VideoFreeze (MedAssociates, # SOF-843). Additional analyses of motor activity during days 1 and 2 were performed with MATLAB (MathWorks, Natick, MA). Chamber floors and walls were cleaned with 70% ethanol between runs, and chambers were wiped with 0.01% acetic acid before placing animals in their chambers.

Approximately 2 hr after the end of Day 2 of the CFC paradigm, mice were injected intraperitoneally with 4-hyroxytamoxifen (at a dose of 25 mg/kg) to permanently induce td-Tomato expression in activated neurons. 4-hydroxytamoxifen (HelloBio, Cat# HB6040) was prepared fresh on the injection day. It was dissolved at 20 mg/mL in 190 proof ethanol (Sigma, Cat # E7148) by vortexing, placing on a bath sonicator for 15 min and vortexing again. This solution was mixed at a 1:1 ratio (v/v) with a 1:4 mixture of castor oil: sunflower seed oil (Sigma, Cat #s 259853 and S5007) to yield a final concentration of 10 mg/ml of 4-hydroxytamoxifen. After vortexing, the ethanol was evaporated by centrifugation under vacuum for 30 min (Centrivap Concentrator, model 78100-00, Labronco, Kansas City, MO).

### Histology and Two-Dimensional Cell Counting

Three days after the tamoxifen injection, mice were injected with ketamine/xylazine (250 mg/kg intraperitoneally) and transcardially perfused. The perfusion process started with 32°C phosphate-buffered saline (PBS) containing procaine hydrochloride (1 g/L; Sigma-Aldrich, St. Louis, MO) and heparin (1 USP unit/mL; Sagent Pharmaceuticals, Schaumburg, IL), continuing for 2 minutes. This was followed by a 2-minute perfusion with 4% paraformaldehyde (PFA; Sigma-Aldrich) in PBS (7.4 pH) at room temperature, and then a 5-minute perfusion with ice-cold 4% PFA in PBS. The brains were then removed, post-fixed in 4% PFA in the dark at 4°C on a rotating shaker for 48 hours, and subsequently cryoprotected in 30% sucrose (w/v in PBS) for another 48 hours. Brains were sectioned at 50 µm thickness in the coronal plane using a cryostat (Model 505E, Microm GmbH, Walldorf, Germany), and stored at −20°C in multi-well plates containing a cryoprotective solution (0.05 M phosphate buffer pH 7.4, 25% glycerol, and 25% ethylene glycol). On average, 40 serial 50 µm coronal sections from saline- and ethanol-exposed mice, containing the limbic memory system (spanning from Bregma −0.35mm to Bregma −2.03mm, as per (Franklin & Paxinos, 2013)), were processed for microscopy. After rinsing with PBS, the sections were mounted on Superfrost Plus microscope slides (Cat #48311-703, VWR, Radnor, PA), covered with glass coverslips, and sealed with DAPI-Fluoromount G (Cat # 0100-20, Southern Biotech, Birmingham, AL).

Fluorescence microscopy was conducted using a Zeiss Axioscan Z1 with a 20X□objective (Preclinical Core, Center for Brain Recovery and Repair, UNM-HSC). Each hemisphere of the brain sections yielded two fluorescence channels: blue for DAPI and red for td-Tomato. Two investigators, blinded to the experimental conditions, independently quantified td-Tomato-positive cell density per mm² in the ATN, CA1 and CA3, dentate gyrus (DG), and RSP using QuPath (Belfast, Ireland). The counts were performed automatically using a single measurement classifier with a fluorescence intensity threshold. The resulting data were averaged between the two investigators.

### QUINT

QUINT is a histological processing workflow (Yates et al., 2019). It comprises several different programs that register brain sections to the Allen Brain Atlas (ABA) (Allen Institute, Seattle, WA) common coordinate framework, automatically count features of interest, and then combines the two to produce reports of how many features of interest are in each brain region. First, we converted images into serially labeled low- and high-resolution PNG files. Low-resolution PNG files were utilized in DeepSlice (Carey et al., 2023) to automatically register sections to the ABA. These were then manually refined in QuickNII (Puchades et al., 2019). Feature quantification utilized the high-resolution PNG files and was accomplished using ilastik (Berg et al., 2019).

Information about feature quantification and information about spatial registration were then combined in Nutil (Groeneboom et al., 2020) which subsequently generated CSV files that were imported and analyzed with MATLAB 2023B (MathWorks, USA). CSV files contained raw counts of Fos from each brain region tested. A list of brain regions between ABA adult mouse slices 59 and 70 was generated. All brain regions that did not fall within that list were excluded. All fiber tracts and ventricles were additionally excluded. Brain regions with subdivisions (such as cortical layers) were consolidated. Forty randomly selected sections from each animal were averaged. All Fos counts had 1 added to their total and they were then log10 transformed. This resulted in an NxM matrix where N represents the total number of brain regions analyzed and M represents the total number of animals analyzed.

### Functional network analysis

The NxM matrix was split according to treatment (TTAE vs Saline) and sex (Male vs Female). A correlation matrix was then computed such that the NxM matrix of Fos counts per animal and brain regions became a NxN matrix of Fos correlations between each brain region. We first plotted the raw matrices and assessed common network features. Next, hierarchical agglomerative clustering with complete linkage was performed to determine functional clusters that emerged in the Fos data. The number of clusters and variation of information was calculated between treatment conditions at each dendrogram cutoff point. The dendrogram cutoff point used for subsequent analyses and plotting was determined by finding the maximum value of variation of information. Next, the participation coefficients and within-module z-scores (WMZs) were calculated and assessed for each analyzed brain region. All analyses were performed in MATLAB 2023B using a mixture of custom routines and the Brain Connectivity Toolbox (Rubinov et al., 2009).

### Statistics

Statistics were performed in MATLAB 2023B. For behavioral data, metrics related to freezing behavior and shock sensitivity were calculated from the raw data. These metrics, in addition to factors for both treatment and sex, were put into a general linear mixed-effects model (GLME). Fixed factors were stepwise added to determine those that significantly improved the overall fit of the GLME. In addition to fixed effects, random effects of litter were added to each GLME if they significantly improved the fit of the model. Models were compared using a theoretical likelihood ratio (LR) test. Additionally, a stepwise linear regression was performed with all 120 brain regions as potential factors using MATLAB’s stepwiseglm function. The model comprised both forward and backward steps. Factors were included if they minimized the Akaike Information Criterion (AIC). Lower AIC values indicate a better fit with less complexity and balances model complexity with performance. Values less than 0 were added in the forward step and values greater than 0.1 were removed in the backward step. At each step, the predictor that produced the lowest AIC was added. For all other statistics where litter was not applicable, distributions were first tested for normality and then either parametric or non-parametric tests were applied. For all statistics, alpha was set to 0.05. Outliers were detected and removed using the Generalized Extreme Studentized Deviate test.

## Results

### TTAE results in less recall following CFC

We first calculated the movement associated with shock times on Day 1 of CFC and no difference emerged in the overall movement data (**Figure 1A**). Shock sensitivity was then calculated as the ratio between the slopes of movement during the first 0.5 s of shock versus the slopes of movement during the 0.5 s following the shock (**Figure 1A a/b**). Mixed effect modeling of random litter effects did not produce a significant increase in model fit (LR Test, LR = 0.059, p = 0.808). A significant main effect of both treatment (ANOVA, F(1,144) = 5.13, p = 0.025) and sex (F(1,144) = 8.777, p = 0.036) as well as a significant interaction (F(1,144) = 6.065, p = 0.015) were observed in shock sensitivity (**Figure 1B**). Post-hoc comparisons revealed a significant difference between saline female and saline male shock sensitivity (Tukey’s HSD, p = 0.011, CI: [-1.544, −0.1433]). The percentage of freezing time was automatically assessed via VideoFreeze (MedAssociates; Vermont, USA). Adding litter as a random effect did not improve model fit (LR Test, LR = 3.185, p = 0.074). Percent freezing time was significantly reduced by both treatment (ANOVA, F(1,156) = 12.741, p < 0.001) and sex (F(1,156) = 8.040, p = 0.005) with female mice and TTAE treated animals freezing less (**Figure 1C**). Planned comparisons revealed significant differences between multiple groups. First, the female saline group froze less than the male saline group (Tukey’s HSD, p = 0.026, CI: [0.647, 14.524]). The female EtOH group froze significantly less than the female saline group (p = 0.002, CI: [2.610, 16.483]), the male saline group (p < 0.001, CI: [7.599, 26.664]), and the male EtOH group (p = 0.026, CI: [0.647, 14.524]). Additionally, the male EtOH group froze less than the male saline group (p = 0.002, CI: [2.610, 16.483]). Together these results suggest that TTAE produces lasting deficits to CFC in a sex specific manner. Last, no relationship between the percentage of freezing time and shock sensitivity was observed for either females or males (**Figure 1D.1,D.2**) suggesting that differences in shock sensitivity were not responsible for differences in freezing behavior. Overall, these findings indicate that TTAE produces deficits in contextual fear memory recall in adult TRAP2;Ai14 mice.

**Figure 1.**
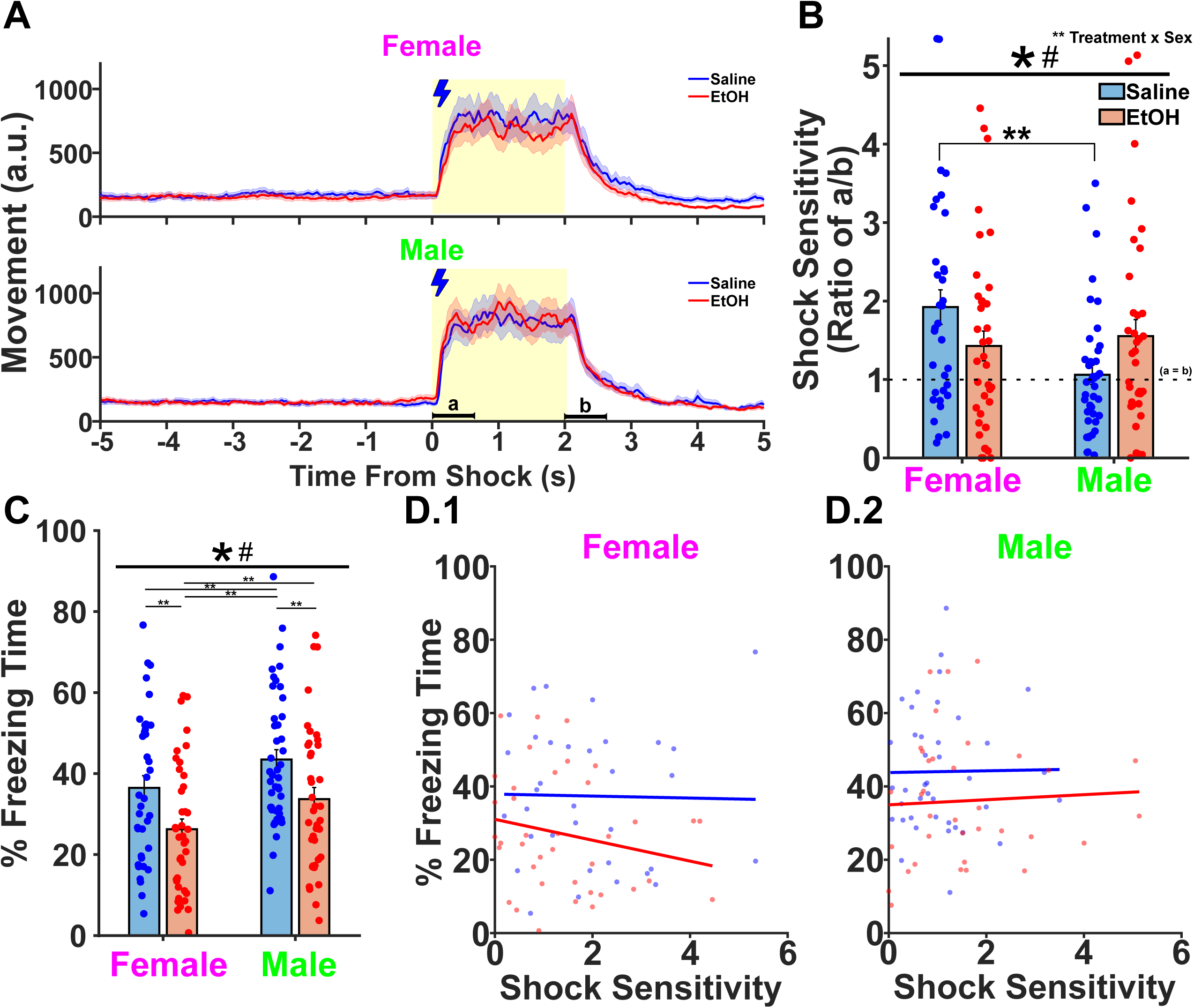
TTAE produces significant decreases in freezing behavior following CFC. **A.** Movement was captured by VideoFreeze and was centered on 4 shock timings. All shocks were pooled together and the mean was taken to assess if there were significant differences in animals’ response to shocks on day 1. The blue lightning bolt and yellow shading indicate time points when the shock was active. Lowercase a and lowercase b indicate the time points wherein our shock sensitivity metric was calculated. **B.** Shock sensitivity, measured as the ratio between a and b, was calculated for each sex and treatment condition per animal. A main effect of both sex (*) and treatment (#) were found with an additional interaction between treatment and sex. Values closer to 1 indicate that movement exclusively happened during the shock period, and a subsequent freeze occurred quickly after the shock ended. Additionally, a post-hoc analysis revealed differences between saline males and saline females (**). **C.** The percentage of the time the animal spent freezing on day 2 was assessed. A main effect of sex and treatment were detected, both lowered the expression of freezing. Significant results of planned comparisons are denoted by (**). **D.1, 2** No correlations between the percentages of time spent freezing or shock sensitivity was detected in females and males.

### The CA1 and CA3 hippocampal regions show an increase in Fos expression following CFC

Computer-assisted quantification of the density of Fos+ neurons was performed by two independent observers. Regions quantified include the ATN (**Figure 2A**), which comprise the anterior dorsal (AD) and anterior ventral nuclei (AV). Hippocampal formation regions quantified include the CA1, CA3, and DG (**Figure 2B**). Last, the RSP was quantified (**Figure 2C**). For the ATN, adding litter as a random effect significantly increased model fit (LR Test, LR = 13.708, p < 0.001), and no main effects were observed in the densities of Fos+ neurons in the ATN (**Figure 2D**). For the CA1, adding litter as a random effect did not significantly improve model fit (LR Test, LR = 2.558, p = 0.110). CA1 Fos+ neuron densities increased as a function of treatment (ANOVA, F(1,122) = 20.593, p < 0.001, **Figure 2E**). Including litter as a random effect significantly increased model fit for CA3 neurons (LR Test, 10.051, p = 0.002) and CA3 Fos+ neuron densities were increased as a function of treatment (F(1,125) = 25.334, p < 0.001, **Figure 2F**). Last, neither the DG or RSP exhibited differences in Fos+ neurons (**Figure 2G-H**) and both model fits were improved by adding litter as a random effect (DG LR Test, LR = 42.734, p < 0.001; RSP LR Test, LR = 29.639, p < 0.001). Overall, we observed that CA1 and CA3 Fos+ neural densities increased as a function of TTAE treatment which may be indicative of TTAE producing deviations away from hippocampal sparse coding.

**Figure 2.**
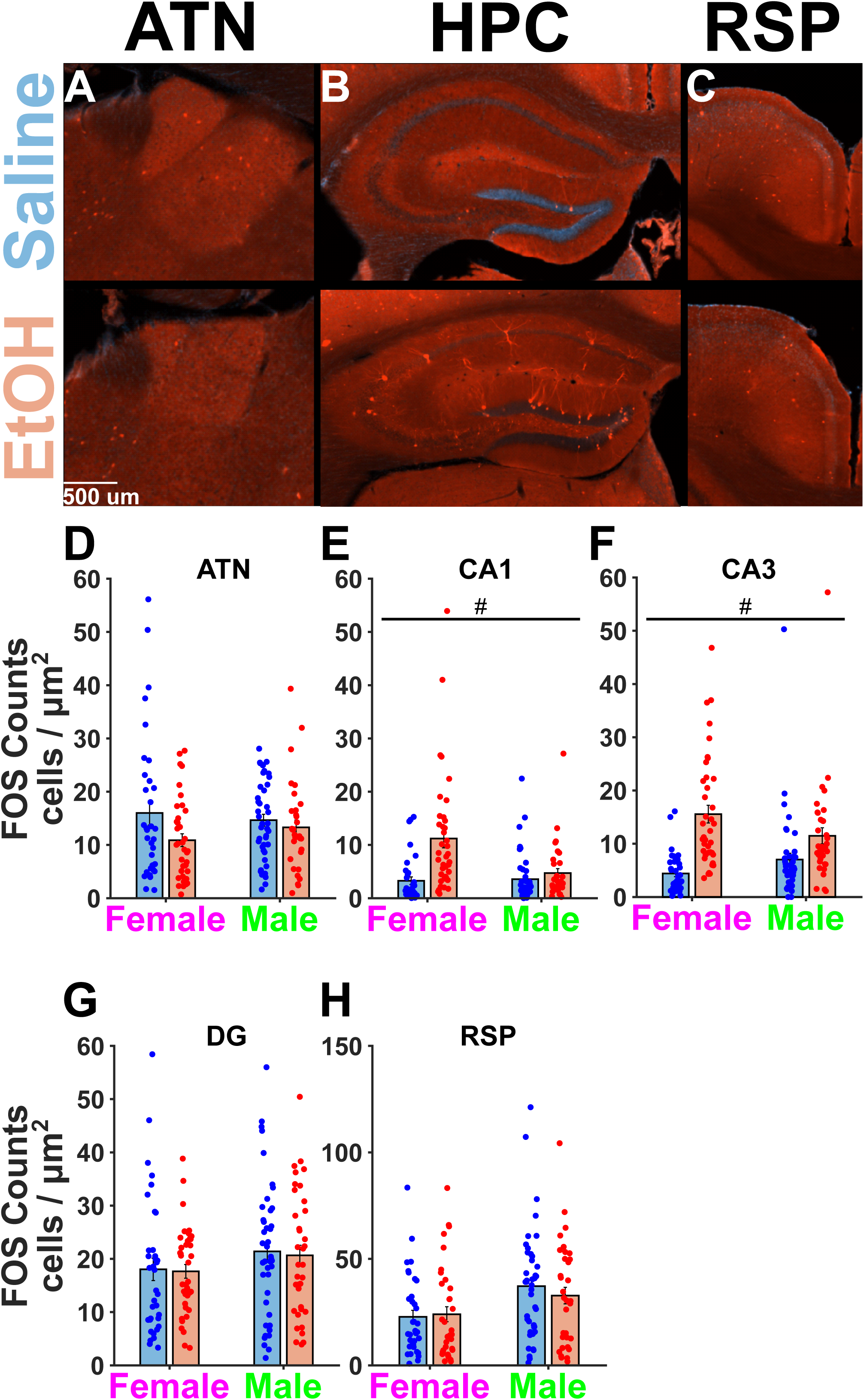
TTAE produces increases in CA1 and CA3 Fos+ neural density following CFC. **A.** Representative images show the ATN. **B.** Representative images show the hippocampal formation including the CA1, CA3, and DG. **C.** Representative images show the RSP. **D.** No main effect of either treatment or sex was detected in the ATN manual counts. **E.** A main effect of treatment, but not sex, was detected in the CA1. **F.** A main effect of treatment, but not sex, was detected in the CA3. **G.** No main effects of either sex or treatment was detected in the DG. **H.** No main effects of either sex or treatment was detected in the RSP.

The two-dimensional Fos counts for female and male ATN showed significant bimodality in the saline treatment condition (**Figure 3A-B**). In order to assuage this potential confound, the ATN nucleus was split into the AD and AV components using an automated cell counting pipeline called QUINT. QUINT was utilized to count Fos expression across the 120 brain regions sampled. In doing so, we obtained counts for the AD and AV. Adding a random effect of litter had no effect on the maximum log likelihood of both the AD and AV GLME models and disallowed comparison between the two models using the LR test as was done above. In the AD, we found a significant decrease in Fos+ neural expression as a function of treatment (F(1,91) = 11.037, p = 0.001, **Figure 3C**). In the AV, we also found a significant decrease in Fos+ neural expression as a function of treatment (F(1,91) = 15.303, p < 0.001, **Figure 3D**). Overall this suggests that individual AD and AV nuclei but not the composite counts are affected following CFC and TTAE.

**Figure 3.**
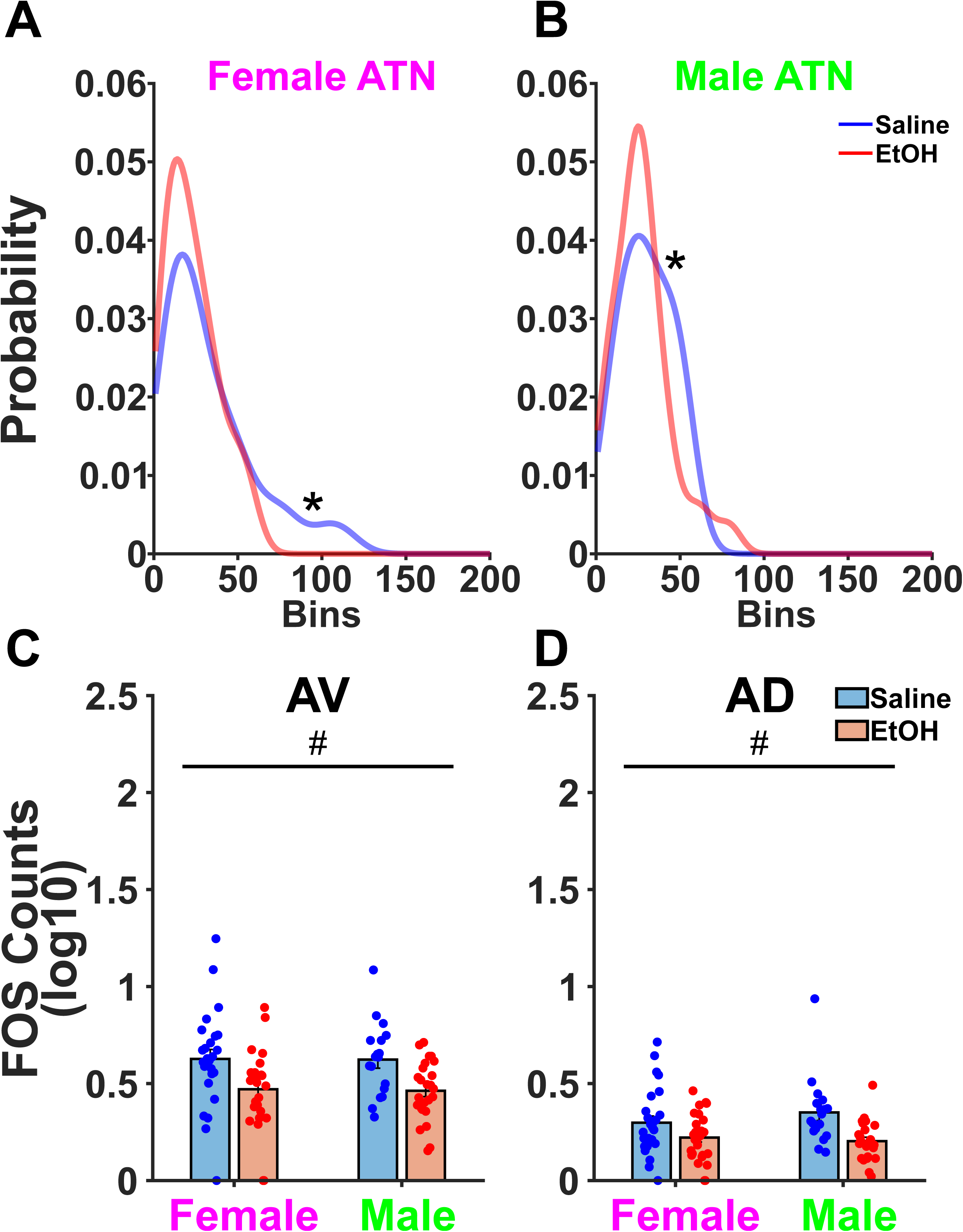
QUINT Further Splits ATN into AD and AV. **A.** A Hartigan’s dip test confirmed bimodality in the manual counts of both female and **B.** male saline groups. **C.** QUINT counts separated the AV and AD that were previously combined as a singular ATN. A significant main effect of treatment was found in the AV wherein TTAE decreased QUINT FOS counts. **D.** A significant main effect of treatment was also found in the AD wherein TTAE decreased QUINT FOS counts.

### Automated counts reveal that CFC motifs of Fos functional connectivity are disrupted following TTAE

Next, we utilized the automated counts to construct an approximation of functional connectivity in the limbic memory system (**Figure 4A**). Brain regions included in our limbic memory circuit included: dorsal premammillary nucleus (PMd), ventral premammillary nucleus (PMv), tuberomammillary nucleus (TM), AD, anteromedial nucleus (AM), AV, subiculum (SUB), DG, CA1-CA3, RSP, and lastly the anterior cingulate cortex (ACA). A common strategy to analyze Fos functional connectivity data is to use agglomerative hierarchical clustering methods. We performed this analysis with complete linkage. Following this analysis, we observed a series of sex differences between the male and female limbic memory circuitry. First, we changed the threshold of the clustering cutoff for several analysis iterations. This revealed that saline males had a sharper decrease (Wilcoxon Rank Sum, z = −4.15, p < 0.001) in the number of clusters formed, whereas females had similar curves (**Figure 4B**). The sharper decrease in indicative of increases in connectivity within the TTAE group resulting in decreases in distances between brain regions. Next, we calculated the variation of information within a sex between the TTAE and saline treatment conditions for each cluster cutoff point tested above. Variation of information was more prominent in males, suggesting a larger difference in brain regions involved in each cluster (**Figure 4C**). We utilized the point of maximal variation of information to determine the cluster cutoff for subsequent analyses. The clustering at this point is shown in **Figure 4D**. Of note, the AV prominently clusters with the DG, SUB, and CA3 following TTAE in both males and females. This suggests that CFC memories engage separate circuits in mice from the saline group vs the TTAE group.

**Figure 4.**
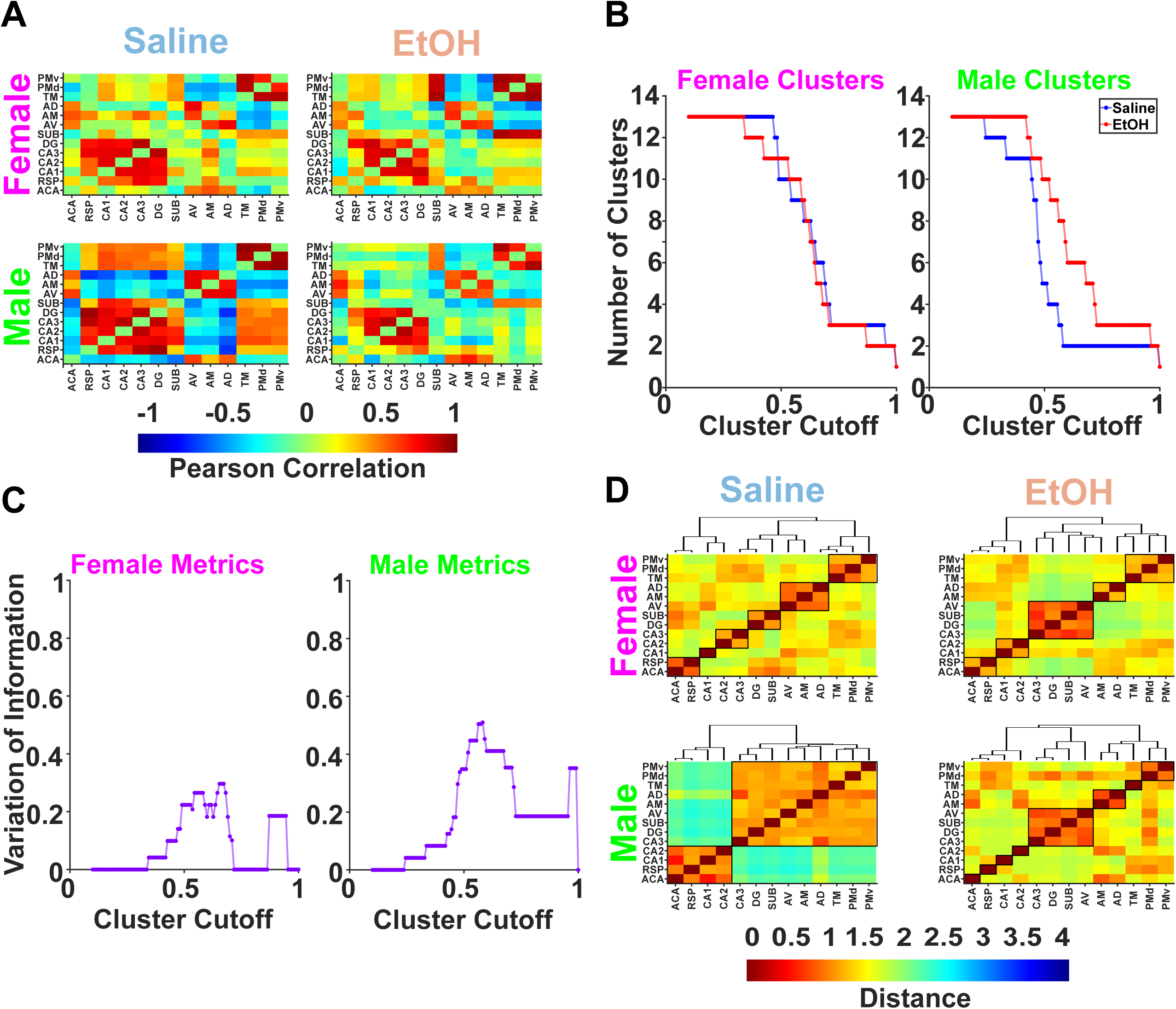
Functional connectivity of the limbic memory system shows alterations following TTAE and CFC. **A.** The Pearson correlations of 13 brain regions part of or connected with the limbic memory system are shown for each sex and treatment condition. Warmer colors indicate higher, positive correlations whereas cooler colors indicate smaller, negative correlations. **B.** Hierarchical agglomerative clustering was performed on the 13 brain regions analyzed. The cluster cutoff refers to the height of the dendrogram wherein 0.5 indicates the halfway point. A rightward shift in EtOH treated animals is observed which suggests differences in clustering emerge in male, EtOH animals. **C.** The variation of information compares the EtOH and Saline clusters for male and female animals to determine at which point they are maximally different. **D.** The point of maximal difference between EtOH and Saline animals is the cluster cutoff point chosen to assign each brain region into its respective clusters. Critically, we see a shift in clustering between the AV and its neighboring anterior thalamic nuclei towards hippocampal sub regions. **Abbreviations.** ACA: Anterior cingulate cortex. RSP: Retrosplenial cortex. DG: Dentate Gyrus. SUB: Subiculum. AV: Anteroventral thalamic nucleus. AM: Anteromedial thalamic nucleus. AD: Anterodorsal thalamic nucleus. TM: Tuberomammillary body. PMd: Dorsal premammillary nucleus. PMv: Ventral premammillary nucleus.

We then considered the functional connectivity of the 120 brain regions analyzed (*Table 1,* **Figure 5A**). The same methodology was used for the larger array as was used in the smaller limbic memory circuitry. In both males and females, a rightward shift was observed in the number of clusters formed following TTAE (**Figure 5B**), but neither reached statistical significance (Wilcoxon Rank Sum, Males: z = −0.645, p = 0.520, Females: z = −1.384, p = 0.166). Additional analysis of the variation of information revealed similar patterns of increasing variation of information between TTAE and saline within each sex (**Figure 5C**). This suggests that although there is only a minor, rightward shift in the number of clusters per cut-off point, the identities of individual brain regions within each of those clusters varies suggesting that different circuits are being recruited during contextual fear memory recall. Last, the point of maximal VI was once again used to determine the cluster cutoff point utilized for the resulting plots and analyses (**Figure 5D**). Overall, linkage distances between nodes decreased in the TTAE condition which is suggestive of hyperconnectivity. Additionally, TTAE appears to produce smaller yet more numerous clusters.

**Figure 5.**
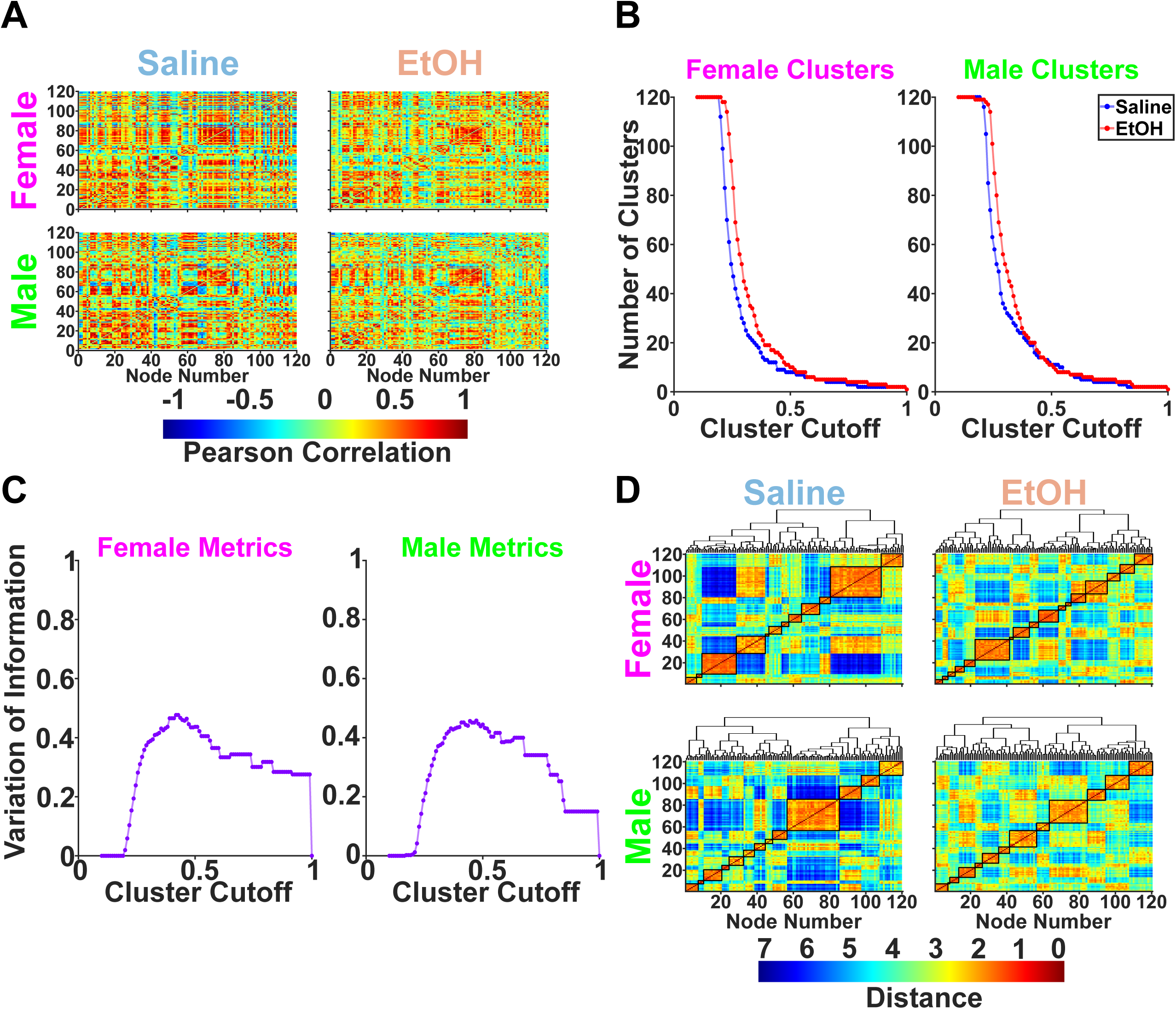
Brainwide hyperconnectivity may emerge following TTAE. **A.** The Pearson correlations of 120 brain regions are shown. Identities of values can be found in *Table 1*. **B.** Hierarchical agglomerative clustering was performed. The cluster cutoff points represent the height of the dendrogram such that 0.5 indicates the number of clusters halfway up the dendrogram. **C.** The variation of information is a metric of how different the cluster identities are between Saline and EtOH conditions. **D.** The point of maximal variation of information is used to assign and plot final clusters. Qualitatively it is observed in the EtOH condition that distances shrink indicated by the presence of warmer colors.

**Table 1.** Full list of 120 brain regions analyzed following QUINT automated counts. All names and abbreviations follow ABA nomenclature.

| Full name | Abbreviation |
| --- | --- |
| Primary motor area | MOp |
| Secondary motor area | MOs |
| Primary somatosensory area | SSp |
| Supplemental somatosensory area | SSs |
| Gustatory areas | GU |
| Visceral area | VISC |
| Dorsal auditory area | AUDd |
| Primary auditory area | AUDp |
| Ventral auditory area | AUDv |
| Anteromedial visual area | VISam |
| **Anterior cingulate area** | **ACA** |
| Agranular insular area | AI |
| Retrosplenial area | RSP |
| Anterior area | VISa |
| Rostrolateral area | VISrl |
| Temporal association areas | TEa |
| Perirhinal area | PERI |
| Ectorhinal area | ECT |
| Olfactory areas | OLF |
| Piriform area | PIR |
| Nucleus of the lateral olfactory tract | NLOT |
| Cortical amygdalar area | COA |
| Piriform-amygdalar area | PAA |
| Field CA1 | CA1 |
| Field CA2 | CA2 |
| Field CA3 | CA3 |
| Dentate gyrus | DG |
| Fasciola cinerea | FC |
| Induseum griseum | IG |
| Entorhinal area | ENT |
| Subiculum | SUB |
| Cortical subplate | CTXsp |
| Clastrum | CLA |
| Endopiriform nucleus | EP |
| Lateral amygdalar nucleus | LA |
| Basolateral amygdalar nucleus | BLA |
| **Basomedial amygdalar nucleus** | **BMA** |
| Posterior amygdalar nucleus | PA |
| Striatum | STR |
| Caudoputamen | CP |
| Fundus of striatum | FS |
| Lateral septal nucleus | LS |
| Septofimbrial nucleus | SF |
| Anterior amygdalar area | AAA |
| Bed nucleus of the accessory olfactory tract | BA |
| Central amygdalar nucleus | CEA |
| Intercalated amygdalar nucleus | IA |
| **Medial amygdalar nucleus** | **MEA** |
| Pallidum | PAL |
| **Globus pallidus** | **GP** |
| Substantia innominata | SI |
| Magnocellular nucleus | MA |
| Triangular nucleus of septum | TRS |
| Bed nuclei of the stria terminalis | BST |
| Ventral anterior-lateral complex of the thalamus | VAL |
| Ventral medial nucleus of the thalamus | VM |
| Ventral posterolateral nucleus of the thalamus | VPL |
| Ventral posteromedial nucleus of the thalamus | VPM |
| Subparafascicular nucleus | SPF |
| **Subparafascicular area** | **SPA** |
| Dorsal part of the lateral geniculate complex | LGd |
| Lateral posterior nucleus of the thalamus | LP |
| Posterior complex of the thalamus | PO |
| Posterior limiting nucleus of the thalamus | POL |
| Ethmoid nucleus of the thalamus | Eth |
| Anteroventral nucleus of thalamus | AV |
| Anteromedial nucleus | AM |
| Anterodorsal nucleus | AD |
| Interanteromedial nucleus of the thalamus | IAM |
| Interanterodorsal nucleus of the thalamus | IAD |
| Lateral dorsal nucleus of thalamus | LD |
| Intermediodorsal nucleus of the thalamus | IMD |
| Mediodorsal nucleus of thalamus | MD |
| Submedial nucleus of the thalamus | SMT |
| Perireunensis nucleus | PR |
| Paraventricular nucleus of the thalamus | PVT |
| Parataenial nucleus | PT |
| Nucleus of reuniens | RE |
| Xiphoid thalamic nucleus | Xi |
| **Rhomboid nucleus** | **RH** |
| **Central medial nucleus of the thalamus** | **CM** |
| Paracentral nucleus | PCN |
| Central lateral nucleus of the thalamus | CL |
| Parafascicular nucleus | PF |
| Reticular nucleus of the thalamus | RT |
| Intergeniculate leaflet of the lateral geniculate complex | IGL |
| Ventral part of the lateral geniculate complex | LGv |
| Subgeniculate nucleus | SubG |
| Medial habenula | MH |
| Lateral habenula | LH |
| **Supraoptic nucleus** | **SO** |
| Accessory supraoptic group | ASO |
| Paraventricular hypothalamic nucleus | PVH |
| Periventricular hypothalamic nucleus | PV |
| Arcuate hypothalamic nucleus | ARH |
| Dorsomedial nucleus of the hypothalamus | DMH |
| Medial preoptic area | MPO |
| Subparaventricular zone | SBPV |
| Suprachiasmatic nucleus | SCH |
| **Subfornical organ** | **SFO** |
| Anterior hypothalamic nucleus | AHN |
| Tuberomammillary nucleus | TM |
| Medial preoptic nucleus | MPN |
| **Dorsal premammillary nucleus** | **PMd** |
| Ventral premammillary nucleus | PMv |
| Ventromedial hypothalamic nucleus | VMH |
| Posterior hypothalamic nucleus | PH |
| **Lateral hypothalamic area** | **LHA** |
| Preparasubthalamic nucleus | PST |
| Parasubthalamic nucleus | PSTN |
| **Perifornical nucleus** | **PeF** |
| Retrochiasmatic area | RCH |
| Subthalamic nucleus | STN |
| Tuberal nucleus | TU |
| Zona incerta | ZI |
| Fields of Forel | FF |
| Median eminence | ME |
| Subcommissural organ | SCO |
| Precommissural nucleus | PRC |
| Anterior pretectal nucleus | APN |

### Exploratory analyses identify brain regions of interest

Utilizing the cluster identities from the previous analysis of 120 brain regions, the WMZ and PC for each brain region was computed. WMZ is a measure of how specifically connected an individual brain region is to its cluster, with large positive values indicating high degrees of within-cluster connectivity and low degrees of out-of-cluster connectivity. No major differences emerged in the overall distribution of WMZs in males and females from the saline and TTAE groups (**Figure 6A**). However, when one looks at the difference between specific regions, some strongly increased in WMZ (**Figure 6B**). All regions where differences exceeded the 95^th^ percentile for males and females were labeled in red. For males, those regions include the arcuate hypothalamic nucleus (ARH), suprachiasmatic nucleus (SCH), intermediodorsal nucleus (IMD), anteromedial visual area (VISam), xiphoid thalamic nucleus (Xi), and the striatum (STR). For females, this included the medial preoptic nucleus (MPN), primary somatosensory area (SSp), medial amygdalar nucleus (MEA), dorsal auditory area (AUDd), globus pallidus (GP), and the ventral posteromedial nucleus (VPM). The PC indicates how much each brain region is connected to both its own cluster as well as other clusters in the network, with higher values indicating more connectivity within and between clusters. The same analysis was repeated for the PCs as was done for the WMZs. The PCs show an increase in females from the TTAE group (Wilcoxon Rank Sum, z = −12.63, p < 0.001) and males (Wilcoxon Rank Sum, z = −3.15, p = 0.002) (**Figure 6C**). Taking the difference between treatment conditions and finding those that are above the 95^th^ percentile shows the brain regions that were most prominently affected. The subparafascicular area (SPA) was affected and increased PC in both sexes (**Figure 6D**). In males, the subthalamic nucleus (STN), lateral septum (LS), TM, posterior amygdalar nucleus (PA), and median eminence (ME) were most affected. In females, the subparafascicular nucleus (SPF), parafascicular nucleus (PF), subgeniculate nucleus (SubG), intergeniculate leaflet of the lateral geniculate complex (IGL), and fields of forel (FF) were most affected.

**Figure 6.**
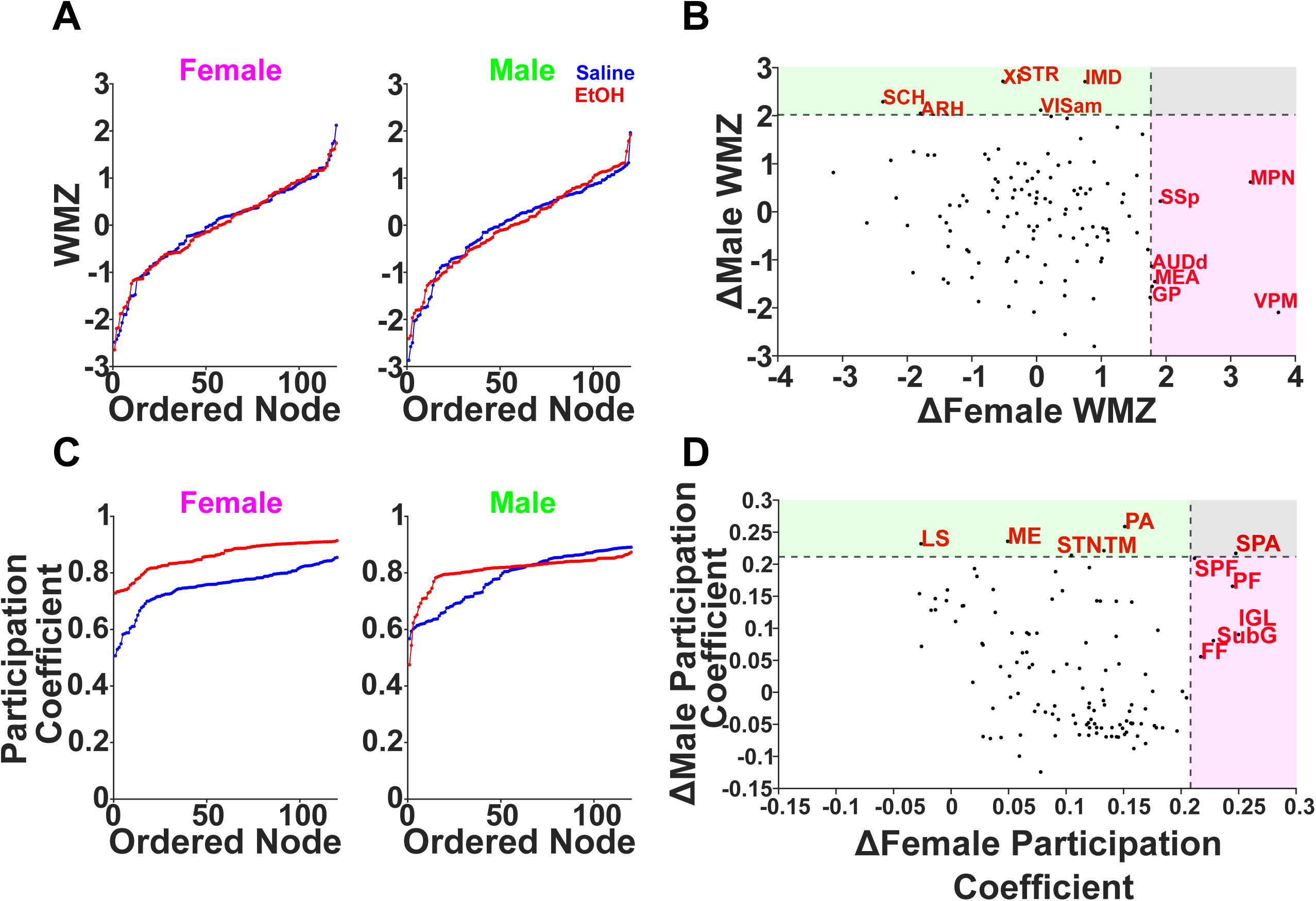
Broad increases in participation coefficients indicate hyperconnectivity with the parafasicular area of the thalamus being most commonly and greatly affected. **A.** Within-Module Z-Scores (WMZs) were calculated for each brain region assessed and showed no overall difference between treatments. **B**. The difference between Saline and EtOH treatment WMZs was calculated for each brain region and sex. The 95th percentile was plotted for each sex with those in green being the highest change in Male WMZ, those in pink being the highest change in Female WMZs and those in gray being high in both. **C**. Participation coefficients were calculated for each sex and treatment condition with the females showing a higher increase following EtOH. **D**. The change in participation coefficient is calculated for each sex. **Abbreviations:** ARH: Arcuate hypothalamic nucleus. MPN: Medial preoptic nucleus. SCH: Suprachiasmatic nucleus. IMD: Intermediodorsal nucleus. VISam: Anteromedial visual area. Xi: Xiphoid thalamic nucleus. STR: Striatum. SSp: Primary somatosensory area. MEA: Medial amygdalar nucleus. AUDd: Dorsal auditory area. GP: Globus pallidus. VPM: Ventral posteromedial nucleus. STN: Subthalamic nucleus. SPF: Subparafascicular nucleus. LS: Lateral septal nucleus. TM: Tuberomamillary nucleus. PA: Posterior amygdalar nucleus. ME: Median eminence. SPA: Subparafascicular area. PF: Parafascicular nucleus. SubG: Subgeniculate nucleus. IGL: Intergeniculate leaflet of the lateral geniculate complex. FF: Fields of Forel.

Next, we sought to determine which, if any, brain regions affected freezing behavior. A stepwise linear model was generated that predicted freezing behavior as a function of sex, treatment, and the Fos expression of 120 brain regions. It iteratively added or removed brain regions as predictor variables in a stepwise fashion until converging on a final set of significant beta values. The model utilized both forward and backward steps utilizing significant p-values (p < 0.05) from sum-of-squared errors in order to add or remove significant predictors. Model coefficient values showed negligible covariance (**Figure 7A**) and had a range of coefficient values (**Figure 7B**). Overall, significant factors included the ACA, basal medial amygdala (BMA), MEA, globus pallidus (GP), rhomboid nucleus (RH), subfornical organ (SFO), PMd, and lateral hypothalamic area (LHA). Significant interactions included a Treatment x BMA interaction as well as SO x PeF interaction. Of particular interest is the Treatment x BMA interaction observed. The BMA appears to have a moderating role in how much freezing behavior is expressed in the TTAE animals (**Figure 7C, D**). The full model and specific coefficient values can be found in *Table 2*.

**Figure 7.**
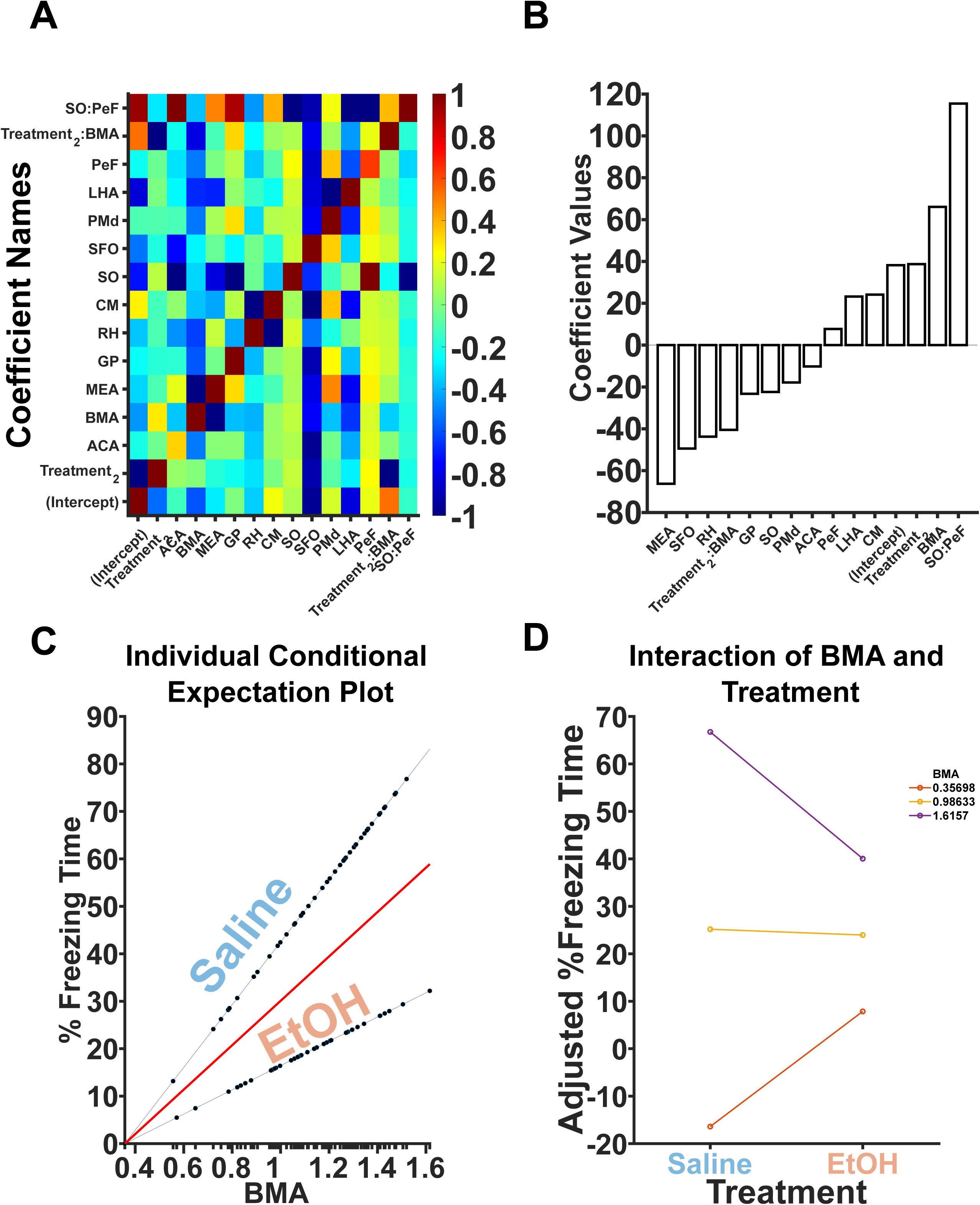
Stepwise linear regression suggests the BMA may be an additional, critical region. **A.** All significant predictor variables and their covariance is shown. Largely, there are no covariant predictors with the exception of interactions. **B.** Coefficient beta values are depicted. Positive values indicate that more Fos+ neurons in that brain region are predictive of increases in freezing. Negative values indicate that more Fos+ neurons in that brain region are predictive of decreases in freezing. **C.** An individual conditional expectation plot is shown depicting the effects of BMA on freezing in both the Saline and EtOH conditions. BMA Fos+ neurons interact with treatment. **D.** BMA interacts with treatment to moderate freezing behavior in EtOH animals.

**Table 2.** Results of stepwise linear-modelling. All significant predictors are marked with bolding.

| Coefficient | Beta | SE | t-value | p-value |
| --- | --- | --- | --- | --- |
| *(Intercept)* | 38.196 | 9.1397 | 4.1792 | **8.43E-05** |
| **Treatment_2** | 38.716 | 11.547 | 3.3528 | **0.0013019** |
| **ACA** | -10.218 | 4.1522 | -2.4609 | **0.016366** |
| **BMA** | 66.06 | 12.389 | 5.3322 | **1.16E-06** |
| **MEA** | -66.27 | 9.2735 | -7.1462 | **7.22E-10** |
| **GP** | -23.283 | 6.0405 | -3.8545 | **0.00025703** |
| **RH** | -43.76 | 17.172 | -2.5484 | **0.013052** |
| CM | 24.156 | 12.454 | 1.9396 | 0.056514 |
| SO | -22.401 | 34.574 | -0.64792 | 0.51919 |
| **SFO** | -49.437 | 19.087 | -2.59 | **0.011698** |
| **PMd** | -17.844 | 5.7683 | -3.0935 | **0.0028553** |
| **LHA** | 23.238 | 11.578 | 2.007 | **0.048666** |
| PeF | 7.7357 | 9.2624 | 0.83518 | 0.4065 |
| **Treatment_2:BMA** | -40.488 | 10.07 | -4.0207 | **0.00014614** |
| **SO:PeF** | 115.46 | 47.227 | 2.4447 | **0.017054** |

## Discussion

TTAE produces significant deficits in CFC. These deficits coincide with increases in CA1 and CA3 Fos expression, and concomitant decreases in AD and AV Fos expression. When Fos is analyzed utilizing measures of functional connectivity, TTAE induces hyperconnectivity across several brain regions resulting in lower likelihoods for those brain regions to form modules with hierarchical agglomerative clustering techniques. Utilizing the clusters determined by hierarchical agglomerative clustering, the WMZ and PC for each brain region was assessed and the difference between treatment conditions was found. PC was significantly affected by TTAE, and the parafasicular area of the thalamus had an increase in the participation coefficient in both males and females. Additionally, stepwise linear regression revealed a significant interaction between TTAE and the BMA Fos counts which may suggest a novel brain region for TTAE’s deleterious effects on CFC.

Bidirectional effects were observed with the CA1 and CA3 exhibiting increased Fos expression following CFC and TTAE while the AD and AV decreased Fos expression following CFC and TTAE. The ATN have been shown to be critical for CFC encoding but not retrieval (de Lima et al., 2017). Additionally, the neural activity of CA1 in conjunction with other regions such as the amygdala is critical for CFC (Jimenez et al., 2020). It is, therefore, a possibility that TTAE disrupts hippocampal-thalamic synchrony during the encoding and/or retrieval of this task and should be explored in future experiments utilizing *in vivo* electrophysiology. Hippocampal subregions are robustly affected by developmental ethanol exposure, particularly when the exposure takes place in the third trimester-equivalent (Coles, 1994). Hippocampus-dependent tasks such as CFC have reliably shown deficits following TTAE (Hamilton et al., 2011; Murawski et al., 2012).

Together, regions of the thalamus and hippocampus comprise the limbic memory system as seen in **Figure 4**. Limbic memory brain regions are disrupted following TTAE treatment due to widespread apoptosis (D. Wozniak et al., 2004). The present study sought to investigate the functional connectivity between these regions utilizing correlations between Fos counts in the disparate limbic memory circuit regions. We saw particularly strong disruptions in the clustering that emerges following TTAE in male subjects. This may reflect differences in how freezing behavior and CFC are encoded between sexes due to a large range of biologically instantiated differences (Keiser et al., 2017; Tronson & Keiser, 2019). A change common to both sexes is the AV becoming more tightly associated with the SUB, DG, and CA3 as opposed to either other anterior thalamic nuclei in females or a wider limbic memory circuit in males. The bidirectional SUB to AV pathway (Shibata, 1993) is critical for spatial memory and is particularly disrupted following the combination of targeted inactivation of the pathway as well as a rotation in an alternating T-maze (Nelson et al., 2020). A working hypothesis is that TTAE produces a deficit in the memory consolidation of CFC wherein more cognitive, spatial effort is required in the TTAE animals prior to exhibiting freezing behavior during day 2. A common theme in the hippocampal literature is that sparse encoding of engrams emerges as a function of learning (Guan et al., 2016; Mao et al., 2017; Wixted et al., 2014) yet our TTAE CA1 and CA3 data violates this assumption suggesting a weakened representation of the environment that must be compensated for via day-of bolstering of the encoding.

Functional connectivity has been shown to be a discriminative measure of FASD (Candelaria-Cook et al., 2023; J. R. Wozniak et al., 2017) which generalizes across both humans and rodents (Rodriguez et al., 2016). Additionally, functional connectivity literature has shown alterations in inter-versus-intra module connectivity in children with FASD versus their typically developing peers (Ware et al., 2021). Interestingly, this aligns well with our differences in PC. Brain networks have been hypothesized to be organized into the so-called “rich-clubs” wherein relatively few brain regions communicate robustly across multiple modules (Heuvel & Sporns, 2011). The PCs increased during TTAE conditions, indicating a departure from this configuration, which may explain aspects of the CFC deficits observed. Alterations in hub properties, such as PC, has been linked with aging (Betzel et al., 2014) and other pathologies such as schizophrenia (Eryilmaz et al., 2022). Overall, alterations in hub properties have been hypothesized to have a significant role in a range of clinical neuropsychiatric conditions (Bullmore & Sporns, 2012; Stam, 2024).

We were surprised that the PC of the SPA was heavily increased in both male and female TTAE mice because this thalamic nucleus is relatively understudied with no alcohol-related studies to date. However, it is apparent that the SPA is involved in the detection of multimodal threats (Kang et al., 2022) with widespread connectivity to cortical and striatal regions (Wang et al., 2006). Additionally, it has been implicated in sex-specific cued-threat learning with females showing greater activation than male counterparts (Du Plessis et al., 2022). Together, this likely implicates the SPA as having a functional role in CFC and it appears to be particularly affected by TTAE. A working hypothesis is that the outsized increase in PC is a compensatory mechanism following TTAE wherein brain regions exhibit hyperconnectivity to maintain function. However, in specific cases such as CFC, the hyperconnectivity has a blocking effect on the encoding and retrieval of fear memories, with a few key nodes having an outsized role, such as the SPA.

Additionally, the BMA emerged as a brain region of interest following its significant interaction with TTAE treatment as predictor of freezing behavior. The BMA has significant roles in the formation of aversive memories via dopaminergic pathways (Zhang et al., 2022) and modulates flight responses in rodents via the superior colliculus (Isa et al., 2021). Given its critical role in fear and emotive memories, it is likely that it is influenced by TTAE and alcohol more broadly. However, to date, only one study focuses on the contributions of prenatal alcohol to amygdala development that includes the BMA and they found significant increases in basolateral amygdala (BLA) neurons but not BMA (Kozanian et al., 2018). Previous studies have shown an increase in BLA excitatory activity following TTAE (Baculis et al., 2015) suggesting that TTAE produces significant functional changes. Future work should include BMA measurements in order to determine if, despite no differences in the number of neurons, the function of the BMA is altered following TTAE.

The present study sought to leverage a large dataset to assess functional connectivity in a TTAE versus saline control model following CFC utilizing Fos-positive neurons. Important limitations in this approach exist. First and foremost, these measurements comprise a single time point. Future studies should investigate how the functional connectivity of these regions changes throughout the entire CFC paradigm, particularly during memory consolidation, using *in vivo* electrophysiology or other measures of neural activity. Additionally, Fos is a measure of excitatory activity and requires a significant amount of activation to transduce. Several neurons may have been just below the required threshold that would not be able to be measured in the present paradigm. Last, the present paradigm utilized a 2D stitching pipeline to produce estimates of 3D volumes from a limited set of brain regions spanning roughly 2 mm posterior from Bregma. Future work should leverage existing brain-clearing technologies in combination with light sheet microscopy in order to obtain larger volumes with higher accuracy.

In conclusion, we found significant decreases in the expression of CFC in TTAE animals. This decrease in freezing behavior was concomitant with increases in CA1 and CA3 Fos expression, and decreases in AV and AD Fos expression. Additionally, we used an automated 2D stitching method to generate a 3D volume of Fos expression for 120 brain regions and found that major alterations in functional connectivity develop following TTAE in response to CFC. A final critical element of these findings was the observed increase in PC, suggesting hyperconnectivity in TTAE brains may directly impact network efficiency, leading to deficits in CFC encoding or retrieval. Last, utilizing a stepwise regression we found a significant interaction between the BMA and TTAE. It is known that TTAE is anxiogenic and produces elevations in excitatory activity in the BLA. However, little is known about the BMA and how TTAE or alcohol more broadly affects its function and structure. Finding common principles by which TTAE affects functional connectivity will aid in developing refined diagnostic criteria for FASD and identifying critical circuitry that can be targeted to ameliorate FASD symptomology. Finding common principles in how functional connectivity measures are affected across tasks, species, and measurement modalities can directly improve clinical outcomes by allowing for cost-effective clinical tools, such as EEG, to be utilized over cost-intensive tools, such as fMRI, in order to improve diagnostic criteria in FASD patients. Together, these results, in conjunction with the present study, suggest that functional connectivity metrics are an indispensable tool in FASD research and identify several novel brain regions that may be affected during CFC following TTAE.

